# High genomic diversity of multi-drug resistant wastewater *Escherichia coli*

**DOI:** 10.1101/215210

**Authors:** Norhan Mahfouz, Serena Caucci, Eric Achatz, Torsten Semmler, Sebastian Guenther, Thomas U. Berendonk, Michael Schroeder

## Abstract

Wastewater treatment plants play an important role in the release of antibiotic resistance into the environment. It has been shown that wastewater contains multi-drug resistant *Escherichia coli*, but information on strain diversity is surprisingly scarce. Here we present an exceptionally large dataset on multidrug resistant *Escherichia coli*, originating from wastewater, over a thousand isolates were phenotypically characterized for twenty antibiotics and for 103 isolates whole genomes were sequenced. To our knowledge this is the first study documenting such a comprehensive diversity of multi-drug resistant *Escherichia coli* in wastewater. The genomic diversity of the isolates was unexpectedly high and contained a high number of resistance and virulence genes. To illustrate the genomic diversity of the isolates we calculated the pan genome of the wastewater *Escherichia coli* and found it to contain over sixteen thousand genes. To analyse this diverse dataset, we devised a computational approach correlating genotypic variation and resistance phenotype, this way we were able to identify not only known, but also candidate resistance genes. Finally, we could verify that the effluent of a wastewater treatment plant will contain multi-drug resistant *Escherichia coli* belonging to clinically important clonal groups.

## Introduction

In 1945, Alexander Fleming, the discoverer of Penicillin, warned of antibiotic resistance. Today, the WHO echoes this warning, calling antibiotic resistance a global threat to human health. Humans are at the center of the modern rise of resistance. The human gut ^1^, clinical samples ^2,3^, soil ^4,5^, and wastewater ^6^ all harbor resistant bacteria and resistance genes. At the heart of modern resistance development is a human-centred network of clinics, industry, private homes, farming, and wastewater. Recent studies suggest that wastewater contains a significant amount of antibiotic resistant *Escherichia coli*, specifically extended-spectrum beta-lactamase-producing *Escherichia coli* ^7^. Particularly, multidrug-resistant (MDR) clones (normally defined as those resistant to three or more drug classes) are of great concern. Past studies have documented the presence of MDR *Escherichia coli* isolates in wastewater on the basis of phenotypic resistance testing ^8^, but a comprehensive analysis of the clonal composition of MDR *Escherichia coli* in wastewater employing whole genome analysis is largely lacking. Therefore the current information on the genomic diversity of antibiotic resistant *Escherichia coli* in wastewater is very limited. Recent metagenomic studies have documented that human-associated bacteria are strongly reduced in the wastewater and its treatment process^9^. To investigate the genomic diversity as well as virulence genes and resistance determinants for wastewater *Escherichia coli*, we proceeded as sketched in Fig. 1: We collected 1178 *Escherichia coli* isolates from a waste treatment plant’s inflow and outflow in the city of Dresden, Germany. We selected 20 antibiotics, which are the most prescribed ones in the area from which the wastewater inflow originates (data provided by the public health insurer AOK). We analyzed the isolates’ resistance to these 20 antibiotics and selected 103 isolates for whole genome sequencing. Our analysis reveals a surprisingly high genomic diversity of MDR *Escherichia coli* in the wastewater with very flexible genomes harboring a high variation of virulence genes and resistance determinants. Using this diversity we developed a computational approach to identify not only known, but also novel candidate resistance genes.

**Figure 1:**
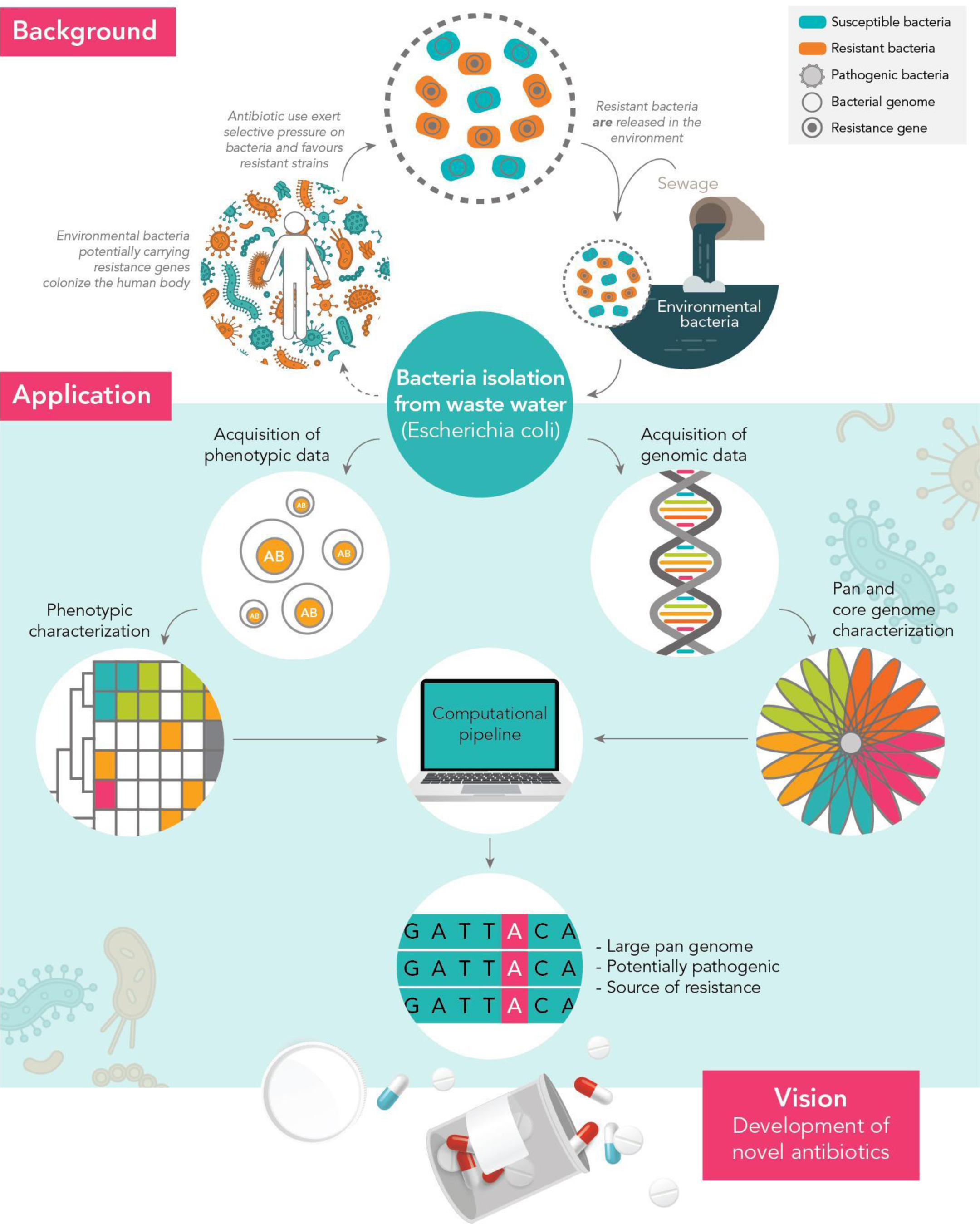
Wastewater plays an important role in antibiotic resistance development. Wastewater *Escherichia coli* isolates are tested for antibiotic resistance and sequenced. Many isolates are multi-drug resistant and potentially pathogenic. Their large pan-genome is a source of potentially novel resistance genes.

## Results

### The wastewater pan-genome

The concept of evolution implies that genomes of organisms of the same species differ. Differences range from small single nucleotide polymorphisms to large genome rearrangements. As a consequence, *Escherichia coli* possesses a core of genes present in all genomes, as well as genes only present in some genomes, or even just in one. The union of all of these genes is called the pan-genome. It is believed, that the *Escherichia coli* core genome comprises around 1400-1500 genes, while the pan-genome may be of infinite size ^10^.

To assess the degree of genomic flexibility of the wastewater isolates, we relate the wastewater pan-genome and the wastewater core genome. At 16582 genes, the wastewater pan-genome is nearly six times larger than the wastewater core genome of 2783 genes, a reservoir of some 14000 genes. Despite this large reservoir, the size difference of nearly 1000 genes between the wastewater *Escherichia coli* core genome and the whole species core genome suggests that the full diversity of *Escherichia coli* is still not covered in our wastewater sample.

The balance between maintaining the core genome and spending energy on acquisition of new genetic material can be captured by the ratio of the core genome size and the average genome size, which is 4700 genes in our sample. This means that only 1400/4700 = 30% of genes in our wastewater *Escherichia coli* are core genes. Most of the non-core genes are very unique and appear only in one or two isolates each. More precisely, 50% of the pan-genome genes appear in only one or two isolates each. This implies that the investigated wastewater *Escherichia coli* are highly individual.

This high diversity is also illustrated in Fig. 2, which compares the wastewater *Escherichia coli* to a clinical dataset of *Escherichia coli*. The figure clearly shows that the *Escherichia coli* of clinical origin are more homogeneous and hence their pan-genome is smaller. In contrast, the diversity of the wastewater *Escherichia coli* match other datasets comprising mixtures of commensal and pathogenic *Escherichia coli*, as well as *Shigella* genomes (see Table 1). This underlines the great diversity of *Escherichia coli* genomes in the wastewater. Interestingly, the variation of the wastewater genomes after the treatment plant was not reduced.

**Figure 2:**
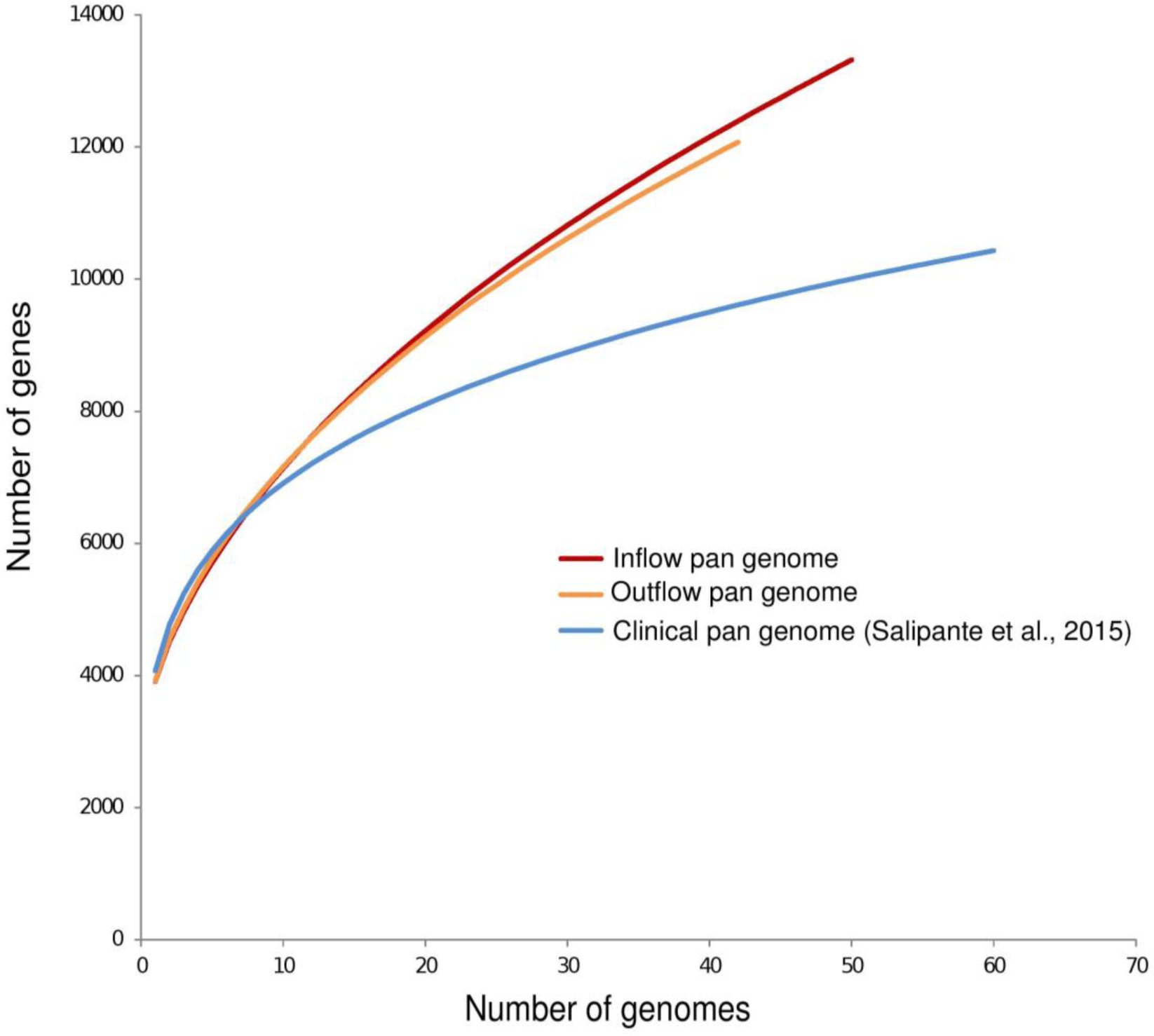
The pan-genome at the outflow has the same size as at the inflow, suggesting that highly flexible *Escherichia coli* emerge from a treatment plant. The wastewater pan-genome is larger than a clinical pan-genome one and of similar size to (see Table 1) highly diverse samples comprising pathogenic, commensal, and lab *Escherichia coli*, as well as *Shigella*.

**Table 1:**
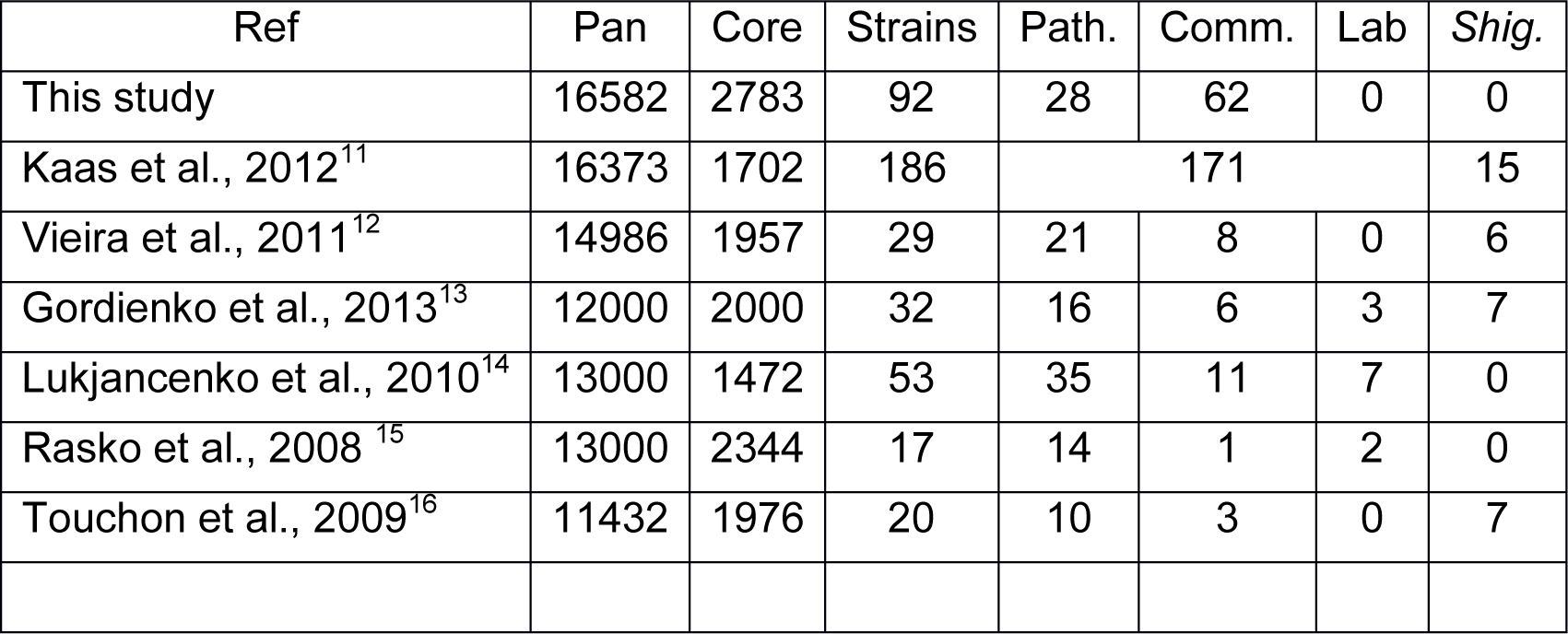
Highly diverse samples comprising pathogenic, commensal, and lab *Escherichia coli*, as well as *Shigella*.

| Ref | Pan | Core | Strains | Path. | Comm. | Lab | Shig. |
| --- | --- | --- | --- | --- | --- | --- | --- |
| This study | 16582 | 2783 | 92 | 28 | 62 | 0 | 0 |
| Kaas et al., 2012<sup>11</sup> | 16373 | 1702 | 186 | 171 |  |  | 15 |
| Vieira et al., 2011<sup>12</sup> | 14986 | 1957 | 29 | 21 | 8 | 0 | 6 |
| Gordienko et al., 2013<sup>13</sup> | 12000 | 2000 | 32 | 16 | 6 | 3 | 7 |
| Lukjancenko et al., 2010<sup>14</sup> | 13000 | 1472 | 53 | 35 | 11 | 7 | 0 |
| Rasko et al., 2008<sup>15</sup> | 13000 | 2344 | 17 | 14 | 1 | 2 | 0 |
| Touchon et al., 2009<sup>16</sup> | 11432 | 1976 | 20 | 10 | 3 | 0 | 7 |

### Resistance genes in the wastewater pan-genome

Wastewater *Escherichia coli* are known to host antibiotic resistance genes. While there are many known resistance genes (see e.g. CARD ^17^), they fall mostly into a few groups, such as beta-lactamases. Here, we seek to confirm and expand the space for candidate resistance genes. Firstly, we measured antibiotic resistance in all 1178 isolates to the 20 antibiotics. As a positive control we included also two antibiotics to which at least clinical *E. coli* are reported to be inherently resistant (kanamycin and cephalotin). Fig. 3 reveals a high degree of resistance and big differences between different antibiotics, including a general trend indicating greater resistance to antibiotics that have been available for longer. Specifically, antibiotics from the 50s and 60s have a significantly different number of resistances than the more recent antibiotics (Welch test, p-value < 0.0025, also significant without including kanamycin and cephalotin). However, there is no significant difference in the number of resistances between isolates from the inflow and the outflow (p-value 0.0001), suggesting that wastewater treatment is not affecting resistance.

**Figure 3:**
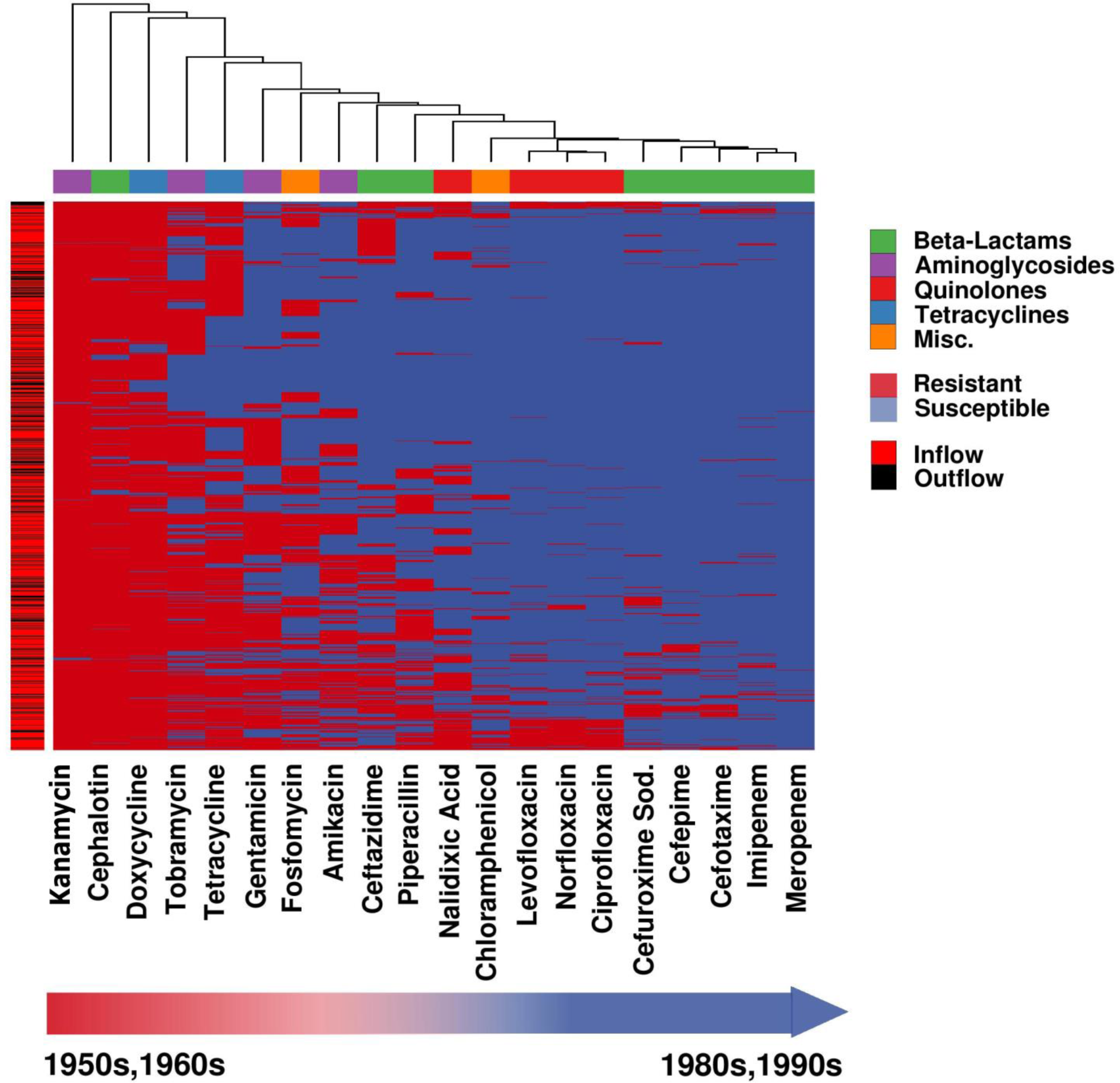
1178 Wastewater *Escherichia coli* isolates are tested for antibiotic resistance to 20 antibiotics. The antibiotics kanamycin and cephalotin were included as a positive control as *E. coli* is reported to be inherently resistant to those antibiotics. Nearly all isolates are multi-drug resistant. Generally, isolates are more susceptible to betalactams and fluoroquinolones than to tetracyclins and aminoglycosides. Surprisingly, the outflow isolates show similar resistance as inflow (p-value 0.0001), suggesting that wastewater treatment is not reducing resistance development.

Next, we correlated the presence of each gene in the sequenced isolates with their phenotypic antibiotic resistance profiles. We excluded meropenem and imipenem, since nearly all isolates are susceptible. For each of the 18 remaining antibiotics, we list the top ten candidate resistance genes in Table 2. These 180 genes comprise 88 unique confirmed genes, including many well-known resistance genes, such as efflux pumps (MT1297 and *emr*E), membrane and transport proteins (*aida-*I*, yia*V*, yij*K*, pit*A*, ics*A, and *pag*N), tetracycline (*tet*A, *tet*R, and *tet*C), chloramphenicol (*cat*), and piperacillin (the beta lactamase *bla*2) resistance genes. However, the 180 genes also comprise a large number of open reading frames encoding hypothetical proteins (41) and genes not yet linked to antibiotic resistance (116). These genes have to be studied further to determine whether they are novel resistance genes or just correlating (e.g. because they are on the same genetic element with a resistance gene). Nearly all of the identified genes are found both in inflow and outflow genomes suggesting that the waste water treatment does not impact on the presence or absence of known and candidate resistance genes.

**Table 2.** Known and candidate resistance genes from correlation of genomes to resistance phenotype. Top 10 genes for 18 antibiotics.

|  | Amikacin | Gentamicin | Kanamycin | Tobramycin | Doxycycline | Tetracycline | Cefepime | Cefotaxime | Ceftazidime | Cefuroxime Sod. | Cephalotin | Piperacillin | Ciprofloxacin | Levofloxacin | Nalidixic Acid | Norfloxacin | Chloramphenicol | Fosfomycin |
| --- | --- | --- | --- | --- | --- | --- | --- | --- | --- | --- | --- | --- | --- | --- | --- | --- | --- | --- |
| 1 | Hypothetical Protein | 4-hydroxyacetophenone monooxygenase **hapE** | Transposase IS200 like protein | Autotransporter precursor **aida-I** | Tetracycline resistance protein, class B **tetA** | Oxygen-dependent choline dehydrogenase **betA** | Ash protein family protein | Hypothetical Protein | cell division protein | Type-1 restriction enzyme R protein **hsdR** | GTPase **era** | Beta-lactamase TEM precursor **bla** | Virulence regulon transcriptional activator **virB** | Transposon Tn10 protein **tetD** | Mercuric resistance operon regulatory protein **merR** | Transposon Tn10 protein **tetD** | Chloramphenicol acetyltransferase **cat** | Invasin |
| 2 | Caudovirales tail fiber assembly protein | Phosphoadenosine phosphosulfate reductases | putative multidrug-efflux transporter/M T1297 | putative protease **yhbU** precursor | Tetracycline repressor protein class B **tetR** | NAD/NADP-dependent betaine aldehyde dehydrogenase **betB** | Fibronectin type III protein | Hypothetical Protein | Plasmid stability protein | Type I restriction enzyme EcoKI M protein **hsdM** | Prophage CP4-57 regulatory protein **alpA** | Transposon Tn3 resolvase **tnpR** | Sporulation initiation inhibitor protein **Soj** | Tetracycline resistance protein, class B **tetA_1** | Mercuric resistance protein **merC** | Tetracycline resistance protein, class B **tetA_1** | Streptomycin 3"-adenylyltransferase **ant1** | Putative DNA-invertase Rac **pinR** |
| 3 | Swarming motility protein **ybiA** | putative multidrug-efflux transporter/M T1297 | Phosphotransferase enzyme family protein | Chaperone protein **dnaK** | Transposon Tn10 TetC protein **tetC** | HTH-type transcriptional regulator **betI** | Transcriptional activator **perC** | Transcriptional activator **perC** | HTH-type transcriptional regulator **cmtR** | **mrr** restriction system protein | Hypothetical Protein | Tyrosine recombinase **xerD** | putative HTH-type transcriptional regulator | Tetracycline repressor protein class B from transposon Tn10 **tetR** | mercuric transport protein **merT** | Tetracycline repressor protein class B from transposon Tn10 **tetR** | Chromosome-partitioning ATPase **soj** | Transcriptional repressor **dicA** |
| 4 | Phospholipase **ytpA** | Phosphotransferase enzyme family protein | Hypothetical Protein | putative ABC transporter ATP-binding protein **yjiK** | HTH-type transcriptional regulator **cmtR** | Tetracycline resistance protein, class B **tetA** | Hypothetical Protein | Hypothetical Protein | Phage-related minor tail protein | Outer membrane protein **lcsA** precursor | Hypothetical Protein | Acetyltransferase (GNAT) family protein | DNA-binding transcriptional regulator **dicC** | Transposon Tn10 protein **tetC** | Mercuric transport protein periplasmic component precursor **merP** | Transposon Tn10 protein **tetC** | **parG** | Hypothetical Protein |
| 5 | Carbonic anhydrase 1 **cynT** | Hypothetical Protein | Streptomycin 3"-adenylyltransferase **ant1** | cell envelope integrity inner membrane protein **tolA** | Tetracycline resistance protein, class C **tetA** | Tetracycline repressor protein class B **tetR** | Hypothetical Protein | Hypothetical Protein | Phage tail protein E | Hypothetical Protein | Hypothetical Protein | Virulence regulon transcriptional activator **virB** | Hypothetical protein | putative HTH-type transcriptional regulator | Anti-adaptor protein **iraM** | CAAX amino terminal protease self-immunity | Hypothetical Protein | Hypothetical Protein |
| 6 | Hypothetical Protein | Hypothetical Protein | Hypothetical Protein | Inner membrane protein **yiaV** precursor | putative inner membrane transporter **yedA** | Transposon Tn10 TetC protein **tetC** | Chromosome partition protein **smc** | Hypothetical Protein | Hypothetical Protein | Fibronectin type III protein | Transposon Tn10 **tetD** protein | Transposase | Lysine--tRNA ligase **lysS** | DNA-binding transcriptional regulator **dicC** | Hypothetical protein | mRNA interferase **pemK** | Hypothetical Protein | Hypothetical Protein |
| 7 | Xanthine dehydrogenase molybdenum-binding subunit **xdhA** | Hypothetical Protein | Zinc-responsive transcriptional regulator | Entericidin B membrane lipoprotein | Tetracycline repressor protein class A from transposon 1721 **tetR** | High-affinity choline transport protein **betT** | Hypothetical Protein | Invasin | Hypothetical Protein | Hypothetical Protein | putative multidrug-efflux transporter/M T1297 | Tetracycline resistance protein, class B **tetA** | Transposon Tn10 protein **tetD** | Hypothetical protein | Mercuric reductase **merA_1** | Antitoxin **pemI** | Acetyltransferase (GNAT) family protein | Molybdenum cofactor biosynthesis protein A |
| 8 | Nicotinate dehydrogenase FAD-subunit **ndhF** | Hypothetical Protein | **merE** protein | Low-affinity inorganic phosphate transporter 1 **pitA** | Hypothetical Protein | Formate dehydrogenase H **fdhF** | Aldehyde-alcohol dehydrogenase **adhE** | Hypothetical Protein | Tyrosine recombinase **xerC** | Hypothetical Protein | Phosphotransferase enzyme family protein | Tetracycline repressor protein class B **tetR** | Tetracycline resistance protein, class B **tetA_1** | CAAX amino terminal protease self-immunity | Hypothetical protein | putative HTH-type transcriptional regulator | putative multidrug-efflux transporter/M T1297 | ATP-dependent zinc metalloprotease **ftsH4** |
| 9 | Nicotinate dehydrogenase small FeS subunit **ndhS** | Phage polarity suppression protein **psu** | Phosphoadenosine phosphosulfate reductases | Methyl-accepting chemotaxis protein II **tar** | Transposon Tn10 **tetD** protein | S-fimbrial protein subunit **sfaH** | Aldehyde-alcohol dehydrogenase **adhE** | Hypothetical Protein | Hypothetical Protein | Hypothetical Protein | Outer membrane protein **pagN** precursor | Transposon Tn10 **tetC** protein | Tetracycline repressor protein class B transposon Tn10 **tetR** | mRNA interferase **pemK** | zinc-responsive transcriptional regulator | DNA-binding transcriptional regulator **dicC** | Phosphotransferase enzyme family protein | Molybdenum cofactor biosynthesis protein A |
| 10 | putative fimbrial-like protein ElfG precursor **elfG** | DNA primase **traC** | Caudovirales tail fiber assembly protein | Leucine-specific-binding protein precursor **livK** | putative multidrug-efflux transporter/M T1297 | Beta-lactamase TEM precursor **bla** | Cob(I)yrinic acid a,c-diamide adenosyltransferase **yvaK** | Type-1 restriction enzyme R protein **hsdR** | Hypothetical Protein | Hypothetical Protein | Tetracycline resistance protein, class B **tetA** | Multidrug transporter **emrE** | Transposon Tn10 protein **TetC** | Antitoxin **PemI** | MerE protein | Caudovirales tail fiber assembly protein | Leucine-specific-binding protein precursor **livK** | Hypothetical Protein |

**Table 3.** Accession numbers of 92 de novo assembled wastewater *Escherichia coli* genomes.

| Bioproject | Biosample | Accession | strain |
| --- | --- | --- | --- |
| PRJNA380388 | SAMN06641941 | NBBP00000000 | Escherichia coli Win2013_WWKa_OUT_3 |
| PRJNA380388 | SAMN06641940 | NBBQ00000000 | Escherichia coli Win2013_WWKa_OUT_29 |
| PRJNA380388 | SAMN06641933 | NBBR00000000 | Escherichia coli Win2013_WWKa_OUT_18 |
| PRJNA380388 | SAMN06641932 | NBBS00000000 | Escherichia coli Win2013_WWKa_OUT_24 |
| PRJNA380388 | SAMN06641931 | NBBT00000000 | Escherichia coli Win2013_WWKa_OUT_1 |
| PRJNA380388 | SAMN06641928 | NBBU00000000 | Escherichia coli Win2013_WWKa_NEU_65 |
| PRJNA380388 | SAMN06641927 | NBBV00000000 | Escherichia coli Win2013_WWKa_NEU_20 |
| PRJNA380388 | SAMN06641926 | NBBW00000000 | Escherichia coli Win2013_WWKa_NEU_60 |
| PRJNA380388 | SAMN06641901 | NBBX00000000 | Escherichia coli Win2013_WWKa_ALT_23 |
| PRJNA380388 | SAMN06641884 | NBBY00000000 | Escherichia coli Win2012_WWKa_OUT_49 |
| PRJNA380388 | SAMN06641883 | NBBZ00000000 | Escherichia coli Win2012_WWKa_OUT_8 |
| PRJNA380388 | SAMN06641882 | NBCA00000000 | Escherichia coli Win2012_WWKa_OUT_34 |
| PRJNA380388 | SAMN06641881 | NBCB00000000 | Escherichia coli Win2012_WWKa_OUT_35 |
| PRJNA380388 | SAMN06641880 | NBCC00000000 | Escherichia coli Win2012_WWKa_OUT_29 |
| PRJNA380388 | SAMN06641879 | NBCD00000000 | Escherichia coli Win2012_WWKa_OUT_26 |
| PRJNA380388 | SAMN06641878 | NBCE00000000 | Escherichia coli Win2012_WWKa_OUT_33 |
| PRJNA380388 | SAMN06641877 | NBCF00000000 | Escherichia coli Win2012_WWKa_OUT_21 |
| PRJNA380388 | SAMN06641876 | NBCG00000000 | Escherichia coli Win2012_WWKa_OUT_2 |
| PRJNA380388 | SAMN06641875 | NBCH00000000 | Escherichia coli Win2012_WWKa_NEU_7 |
| PRJNA380388 | SAMN06641874 | NBCI00000000 | Escherichia coli Win2012_WWKa_OUT_14 |
| PRJNA380388 | SAMN06641873 | NBCJ00000000 | Escherichia coli Win2012_WWKa_NEU_51 |
| PRJNA380388 | SAMN06641872 | NBCK00000000 | Escherichia coli Win2012_WWKa_NEU_31 |
| PRJNA380388 | SAMN06641871 | NBCQ00000000 | Escherichia coli Win2012_WWKa_NEU_37 |
| PRJNA380388 | SAMN06641870 | NBCR00000000 | Escherichia coli Win2012_WWKa_NEU_16 |
| PRJNA380388 | SAMN06641869 | NBCS00000000 | Escherichia coli Win2012_WWKa_NEU_19 |
| PRJNA380388 | SAMN06641868 | NBCT00000000 | Escherichia coli Win2012_WWKa_NEU_12 |
| PRJNA380388 | SAMN06641867 | NBCU00000000 | Escherichia coli Win2012_WWKa_ALT_65 |
| PRJNA380388 | SAMN06641866 | NBCV00000000 | Escherichia coli Win2012_WWKa_NEU_1 |
| PRJNA380388 | SAMN06641865 | NBCW00000000 | Escherichia coli Win2012_WWKa_ALT_49 |
| PRJNA380388 | SAMN06641864 | NBCX00000000 | Escherichia coli Win2012_WWKa_ALT_54 |
| PRJNA380388 | SAMN06641863 | NBCY00000000 | Escherichia coli Sum2013_WWKa_OUT_5 |
| PRJNA380388 | SAMN06641862 | NBCZ00000000 | Escherichia coli Sum2013_WWKa_OUT_39 |
| PRJNA380388 | SAMN06641861 | NBDA00000000 | Escherichia coli Sum2013_WWKa_OUT_49 |
| PRJNA380388 | SAMN06641860 | NBDB00000000 | Escherichia coli Sum2013_WWKa_OUT_3 |
| PRJNA380388 | SAMN06641859 | NBDC00000000 | Escherichia coli Sum2013_WWKa_OUT_31 |
| PRJNA380388 | SAMN06641858 | NBDD00000000 | Escherichia coli Sum2013_WWKa_OUT_2 |
| PRJNA380388 | SAMN06641857 | NBDE00000000 | Escherichia coli Sum2013_WWKa_OUT_21 |
| PRJNA380388 | SAMN06641856 | NBDF00000000 | Escherichia coli Sum2013_WWKa_NEU_53 |
| PRJNA380388 | SAMN06641855 | NBDG00000000 | Escherichia coli Sum2013_WWKa_NEU_46 |
| PRJNA380388 | SAMN06641854 | NBDH00000000 | Escherichia coli Sum2013_WWKa_NEU_39 |
| PRJNA380388 | SAMN06641853 | NBDI00000000 | Escherichia coli Sum2013_WWKa_ALT_44 |
| PRJNA380388 | SAMN06641852 | NBDJ00000000 | Escherichia coli Sum2013_WWKa_NEU_29 |
| PRJNA380388 | SAMN06641851 | NBDK00000000 | Escherichia coli Spr2013_WWKa_OUT_27 |
| PRJNA380388 | SAMN06641844 | NBDL00000000 | Escherichia coli Sum2013_WWKa_ALT_41 |
| PRJNA380388 | SAMN06641843 | NBDM00000000 | Escherichia coli Sum2013_WWKa_ALT_27 |
| PRJNA380388 | SAMN06641842 | NBDN00000000 | Escherichia coli Spr2013_WWKa_OUT_56 |
| PRJNA380388 | SAMN06641841 | NBDO00000000 | Escherichia coli Sum2013_WWKa_ALT_20 |
| PRJNA380388 | SAMN06641840 | NBJM00000000 | Escherichia coli Spr2013_WWKa_OUT_5 |
| PRJNA380388 | SAMN06641839 | NBJN00000000 | Escherichia coli Spr2013_WWKa_OUT_55 |
| PRJNA380388 | SAMN06641838 | NBJO00000000 | Escherichia coli Spr2013_WWKa_OUT_32 |
| PRJNA380388 | SAMN06641837 | NBJP00000000 | Escherichia coli Spr2013_WWKa_OUT_45 |
| PRJNA380388 | SAMN06641822 | NBJQ00000000 | Escherichia coli Spr2013_WWKa_OUT_15 |
| PRJNA380388 | SAMN06641821 | NBJR00000000 | Escherichia coli Spr2013_WWKa_OUT_29 |
| PRJNA380388 | SAMN06641820 | NBJS00000000 | Escherichia coli Spr2013_WWKa_NEU_6 |
| PRJNA380388 | SAMN06641819 | NBJT00000000 | Escherichia coli Spr2013_WWKa_OUT_11 |
| PRJNA380388 | SAMN06641818 | NBJU00000000 | Escherichia coli Spr2013_WWKa_NEU_15 |
| PRJNA380388 | SAMN06641817 | NBJV00000000 | Escherichia coli Spr2013_WWKa_NEU_37 |
| PRJNA380388 | SAMN06641816 | NBJW00000000 | Escherichia coli Spr2013_WWKa_ALT_63 |
| PRJNA380388 | SAMN06641815 | NBJX00000000 | Escherichia coli Spr2013_WWKa_ALT_71 |
| PRJNA380388 | SAMN06641814 | NBJY00000000 | Escherichia coli Spr2013_WWKa_ALT_51 |
| PRJNA380388 | SAMN06641813 | NBJZ00000000 | Escherichia coli Spr2013_WWKa_ALT_55 |
| PRJNA380388 | SAMN06641812 | NBKA00000000 | Escherichia coli Spr2013_WWKa_ALT_43 |
| PRJNA380388 | SAMN06641811 | NBKB00000000 | Escherichia coli Spr2013_WWKa_ALT_27 |
| PRJNA380388 | SAMN06641810 | NBKC00000000 | Escherichia coli Spr2013_WWKa_ALT_41 |
| PRJNA380388 | SAMN06641809 | NBKD00000000 | Escherichia coli Spr2012_WWKa_OUT_37 |
| PRJNA380388 | SAMN06641808 | NBKE00000000 | Escherichia coli Spr2012_WWKa_OUT_54 |
| PRJNA380388 | SAMN06641807 | NBKF00000000 | Escherichia coli Spr2012_WWKa_OUT_25 |
| PRJNA380388 | SAMN06641806 | NBKG00000000 | Escherichia coli Spr2012_WWKa_OUT_3 |
| PRJNA380388 | SAMN06641805 | NBKH00000000 | Escherichia coli Spr2012_WWKa_OUT_16 |
| PRJNA380388 | SAMN06641804 | NBKI00000000 | Escherichia coli Spr2012_WWKa_OUT_13 |
| PRJNA380388 | SAMN06641803 | NBKJ00000000 | Escherichia coli Spr2012_WWKa_NEU_74 |
| PRJNA380388 | SAMN06641802 | NBKK00000000 | Escherichia coli Spr2012_WWKa_OUT_12 |
| PRJNA380388 | SAMN06641801 | NBKL00000000 | Escherichia coli Spr2012_WWKa_NEU_31 |
| PRJNA380388 | SAMN06641800 | NBKM00000000 | Escherichia coli Spr2012_WWKa_NEU_51 |
| PRJNA380388 | SAMN06641799 | NBKN00000000 | Escherichia coli Spr2012_WWKa_NEU_24 |
| PRJNA380388 | SAMN06641798 | NBKO00000000 | Escherichia coli Spr2012_WWKa_ALT_27 |
| PRJNA380388 | SAMN06641797 | NBKP00000000 | Escherichia coli Spr2012_WWKa_ALT_35 |
| PRJNA380388 | SAMN06641796 | NBKQ00000000 | Escherichia coli Aut2013_WWKa_OUT_3 |
| PRJNA380388 | SAMN06641793 | NBKR00000000 | Escherichia coli Aut2013_WWKa_OUT_10 |
| PRJNA380388 | SAMN06641792 | NBKS00000000 | Escherichia coli Aut2013_WWKa_OUT_20 |
| PRJNA380388 | SAMN06641791 | NBKT00000000 | Escherichia coli Aut2013_WWKa_NEU_51 |
| PRJNA380388 | SAMN06641789 | NBKU00000000 | Escherichia coli Aut2013_WWKa_NEU_53 |
| PRJNA380388 | SAMN06641788 | NBKV00000000 | Escherichia coli Aut2013_WWKa_NEU_44 |
| PRJNA380388 | SAMN06641786 | NBKW00000000 | Escherichia coli Aut2013_WWKa_ALT_65 |
| PRJNA380388 | SAMN06641785 | NBKX00000000 | Escherichia coli Aut2013_WWKa_NEU_28 |
| PRJNA380388 | SAMN06641784 | NBKY00000000 | Escherichia coli Aut2013_WWKa_ALT_59 |
| PRJNA380388 | SAMN06641782 | NBKZ00000000 | Escherichia coli Aut2013_WWKa_ALT_48 |
| PRJNA380388 | SAMN06641780 | NBLA00000000 | Escherichia coli Aut2013_WWKa_ALT_45 |
| PRJNA380388 | SAMN06641779 | NBLB00000000 | Escherichia coli Aut2013_WWKa_ALT_30 |
| PRJNA380388 | SAMN06641778 | NBLC00000000 | Escherichia coli Aut2013_WWKa_ALT_17 |
| PRJNA380388 | SAMN06641777 | NBLD00000000 | Escherichia coli Aut2013_WWKa_ALT_13 |
| PRJNA380388 | SAMN06670745 | NBNO00000000 | Escherichia coli Win2012_WWKa_OUT_19 |

### Virulence genes

Generally, *Escherichia coli* strains exhibit great variation. Many exist as harmless commensals in the human gut, but some are classified as intra-(InPEC) or extra-intestinal pathogenic *Escherichia coli* (ExPEC ^18^). Based on their virulence genes profile the pathogenic potential of Escherichia coli isolates can be determined^7^. The sequenced isolates contain some 700 of the 2000 *Escherichia coli* virulence factors in the virulence factor database ^19^, averaging to 153 and to 155 virulence factors per isolate for inflow and outflow, respectively. Hence, there is no significant difference (Welch test, CI 95%) between inflow and outflow. In particular, we found combinations of virulence factors for 16 isolates (see methods), which are indicative of ExPEC. Eight of these 16 isolates were obtained from the outflow of the treatment plant (see Fig. 4).

**Figure 4:**
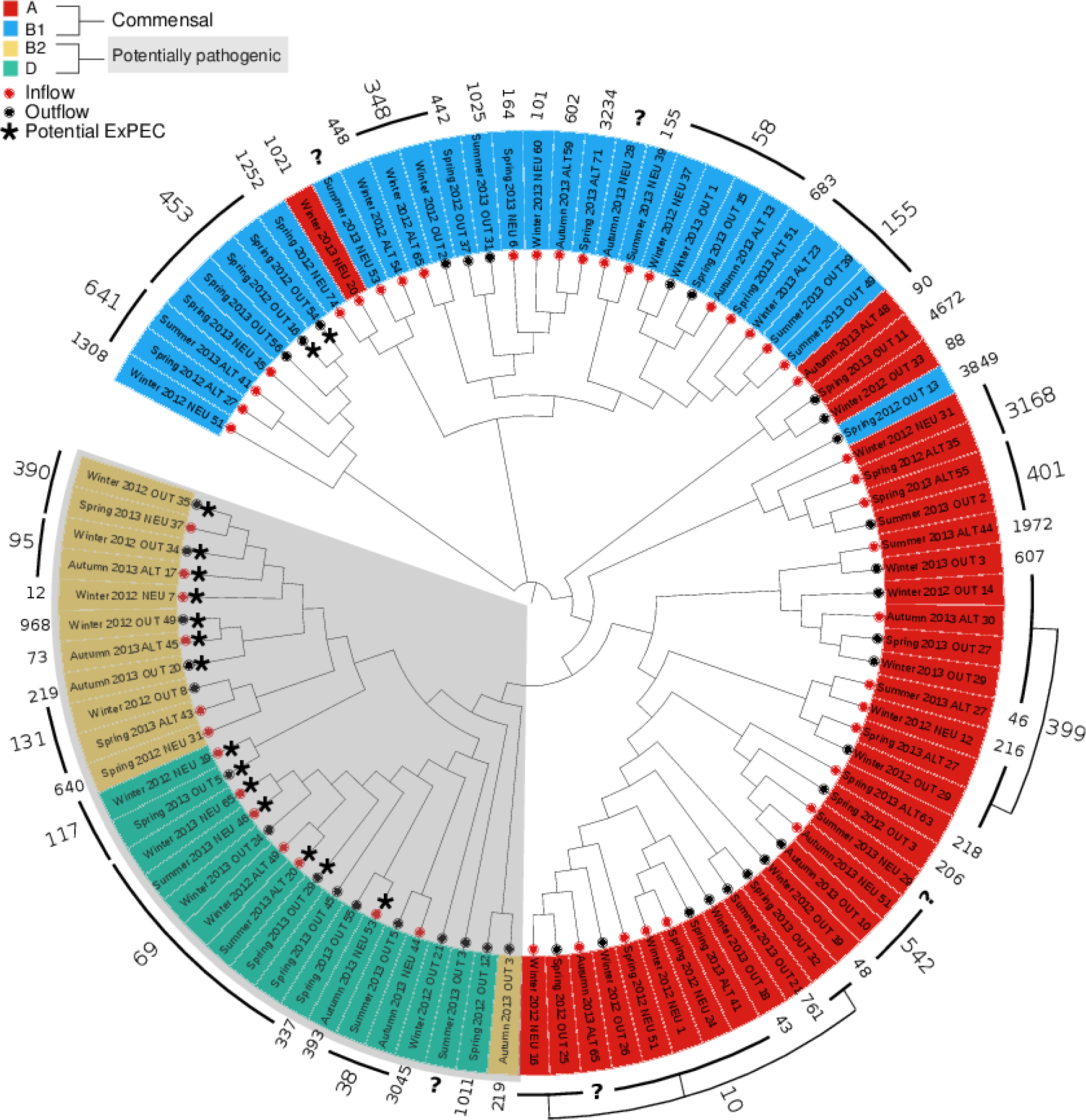
Phylogeny and pathogenic potential of wastewater *Escherichia coli*. Phylogenetic tree, multi-locus sequence types, and phylogroups of 92 sequenced wastewater *Escherichia coli* isolates reveal 16 potential ExPEC isolates (marked with a black star) in phylogroups B2 (yellow) and D (green), which are associated with pathogenicity. Half of the potentially pathogenic isolates stem from the outflow of the treatment plant.

Besides the presence of known virulence factors, the pathogenic potential can be assessed using genotyping with multi-locus sequence types ^20^ and phylogroups ^21^. Broadly, *Escherichia coli* has, among other s, four phylogroups, A, B1, B2 and D. Commensal *Escherichia coli* fall mostly into groups A and B1 and ExPEC into B2 and D ^21^. Fig. 4 shows a phylogenetic tree of the sequenced wastewater *Escherichia coli* isolates along with the commensal phylogroups A (red) and B1 (blue) and the pathogenicity-associated groups B2 (yellow) and D (green), as well as the finer-grained multi-locus sequence types. The tree is based on genomic variations compared to the reference genome of *Escherichia coli* K12 MG1655. Fig. 4 reveals that nearly one third of isolates belong to group B2 and D, in which ExPEC are usually found. In particular, B2 and D include 14 of the 16 potential ExPEC isolates. Remarkably, half of the B2 and D isolates are from the wastewater treatment plant's outflow.

## Discussion

### Pan and core genome

It is well known that wastewater treatment reduces the bacterial abundance, in addition a recent metagenomic study has shown that the bacterial community in wastewater is very different to the human gut community and that the number of detected genera is reduced in the wastewater^9^. Consequently, our expectation was that the genomic diversity of *Escherichia coli* should be reduced. We were very surprised to find an unexpectedly high genomic diversity, which is illustrated in the large pangenome. A possible explanation for this high genomic diversity is that the *Escherichia coli* cells within the wastewater originate not only from human faeces, but also from a multitude of different animal faeces collected via the surface runoff into the sewers. This would also explain why the pangenome of the wastewater *Escherichia coli* is considerably larger than the clinical pangenome reported by Land et al.^22^. Generally, many authors have pointed out that *Escherichia coli* has a large and flexible pan genome. Lapierre *et al*. argue that *Escherichia coli* appears to have unlimited ability to absorb genetic material and hence its pan genome is open ^10^. In a recent study comprising over 2000 genomes Land *et al*. put this into numbers and arrive at a pan genome of 60000-89000 gene families for over 2000 sequenced *Escherichia coli* genomes ^22^. The study by Land *et al*. (24) is based on clinical isolates, in contrast our study is the first, which has calculated the pangenome of *Escherichia coli* for wastewater. Interestingly, our results seem to be in concordance and suggest that within our study we still have not reached the saturation of the detected diversity (Fig. 2), indicating that the full genomic diversity of *Escherichia coli* in the wastewater is probably even larger than what we report here. Worryingly, this is also reflected in a high diversity of resistance and virulence genes. This documents that the wastewater contains a significant amount of multi-drug resistant (MDR) *Escherichia coli*, which also carry a suit of virulence genes suggesting that some of those MDR have a pathogenic potential. Furthermore, we did not find a significant difference in genomic diversity between inflow and outflow of the wastewater treatment plant, suggesting that selection against genome diversity and resistance determinants does not seem to occur.

### Pathogenic potential and resistance

Resistant bacteria may or may not be pathogenic. While ultimate proof for pathogenicity can only be obtained from in vivo studies, we wanted to understand the pathogenic potential of the isolates by analysing the genome for suitable markers. Here we chose to consider three independent approaches: classification by phylogenetic groups, by multi-locus sequence tags, and by identification of specific virulence factors (see methods). While the three approaches showed consistent results, they are by no means proof for pathogenicity, since there can be exceptions to these classification rules. As an example, consider the strain ED1a (O81), which was isolated from a healthy man, but belongs to the phylogenetic group B2 ^16^. Similarly, pathogenicity may not only arise from the acquisition of genes, but also from the loss ^23^.

Regarding resistance there are similar confounding factors. *Escherichia coli* is inherently resistant to kanamycin and cephalotin, which is also clearly shown in Fig. 3. This supports the notion that, generally antibiotic resistance is ancient ^24^ and naturally occurring in the environment. Nonetheless, there are pronounced differences between pristine and human environments ^25^. This is also supported by Fig. 3, which shows that antibiotics introduced in the 60s have more resistances than those introduced later (p-value < 0.0025), which suggests, that the naturally occurring resistances do not play a major role in the emergence of observed resistances.

### From clinic to river

We have shown that there are *Escherichia coli* at the wastewater outflow, which are multi-drug resistant and have pathogenic potential. But are they abundant enough to have an impact in the aquatic system they are released into? They do. The percentage of possibly pathogenic *Escherichia coli* in the outflow is considerable and may correspond to a large absolute amount. If an average of 100 *Escherichia coli* colony forming units (CFU) are released per ml, then 10^13^ CFUs per day are released (assuming a release of 10^5^ m^3^ per day). This is in accordance with Manaia *et al*., who showed that 10^10^-10^14^ CFU of ciprofloxacin-resistant bacteria are released by a mid-sized wastewater treatment plant ^26^. Supporting these results, a study in a Japanese river shows the presence of pathogenic *Escherichia coli* ^27^. In this study they sequenced over 500 samples from the Yamato river and most of their prevalent multi-drug resistant and clinical strains are also present in our samples. In a related study, Czekalski *et al*. found that particle-associated wastewater bacteria are the responsible source for antibiotic resistance genes in the sediments of lake Geneva in Switzerland ^28^. Assuming that the river Elbe is comparable to these aquatic systems, it suggests, that the urban environment (including clinics) and river are connected with wastewater treatment plants in between.

### Composition of phylogroups

It is interesting to compare the breakdown into phylogenetic groups of wastewater *Escherichia coli* to compare samples from human and animal environments. It is, e.g., known that the phylogenetic group B2 is more abundant among commensal *Escherichia coli* from human faeces (43%) than from farm animals (11%) ^29^. Therefore, the composition of wastewater *Escherichia coli* as shown in Fig. 4 resembles commensal *Escherichia coli* from farm animals more closely. Similarly, Tenaillon *et al*. find that groups A and B1 make up one third in human faeces ^29^, whereas we find two thirds. This suggests that animal feces play an important role for resistance also of urban wastewater treatment plants and this is probably part of the explanation for the high observed genomic diversity.

### Random sampling and novel resistance mechanisms

The initial 1178 isolates were sampled randomly over different times of the year, from two different inflows and the outflow of the wastewater treatment plant. In contrast, the 103 sequenced isolates were chosen in such way that all of the phenotypes encountered were represented (see methods). Within a phenotype group isolates were chosen randomly. This random, but representative choice and the subsequent link from genotype to phenotype is an example of high-throughput hypothesis-free analysis. And although, there was no pre-defined resistance mechanism, which we aimed to hit, many of the well-known resistance genes were ranked high. This supports the hope that high-throughput, hypothesis-free methods such as deep sequencing will help to uncover novel resistance mechanisms and in particular that some of the candidate resistance genes will prove to have a causal link to resistance. The results show that the here outlined computational approach to correlate genomic and phenotypic information for wastewater *Escherichia coli* significantly assists to identify a larger part of the existing resistome of *Escherichia coli*.

### Conclusion

Overall, we have shown for the first time that *Escherichia coli* isolates from wastewater have a surprisingly large pan-genome, which harbors virulence genes, known and novel candidate resistance genes. We developed a computational approach based on genomic and phenotypic correlation for *Escherichia coli* and show that applying this to wastewater will discover novel parts of the resistome in *Escherichia coli*. Finally, together with the estimates on absolute *Escherichia coli* abundance, we could demonstrate that there is a considerable pathogenic potential in the outflow of a wastewater treatment plant. Using *Escherichia coli* as an example, this study demonstrates the importance of investigating wastewater with modern bioinformatics and strain specific genomic analysis in order to estimate the extent of genomic variation and resistance determinants for bacteria with clinical relevance present in the environment.

## Methods

### Collection

1178 samples were collected from the municipal wastewater treatment plant Dresden, Germany. Samples were collected on 11/4/2012 (Spring 2012), 30/7/2012 (Summer 2012), 21/1/2013 (Winter 2012), 27/3/2013 (Spring 2013), 6/8/2013 (Summer 2013), 14/10/2013 (Autumn 2013), and 17/12/2013 (Winter 2013). Samples were collected either at the outflow (OUT) or at one of two inflow locations (Altstadt ALT and Neutstadt NEU), representing the area south and north of the river Elbe).

### Isolation

*Escherichia coli* and total coliforms bacteria were enumerated via serial fold dilution plating of the original wastewater (triplicate samples). Wastewaters were diluted in double distilled water, until the enumeration of bacterial colonies was possible. *Escherichia coli* and coliform counts were always performed in triplicates. The *Escherichia coli* colonies were selected and picked after overnight growth at 37°C on a selective chromogenic media (OXOID Brilliance *Escherichia coli*/Coliform Selective Agar, Basingstoke, England). To minimize the risk of colony contamination, picked colonies were spiked a second time on the same selective media and pure single colonies were grown overnight on LB media at 37°C and stored on glycerol stock at −80° C. **Resistance phenotyping.** Antibiotic resistance phenotypes were determined by the agar diffusion method using 20 antibiotic discs (OXOID, England) according to EUCAST (or CLSI when EUCAST was not available) ^7,8^. The selected drugs belong to the most commonly prescribed antibiotics for diseases caused by bacteria according to the German health insurance AOK Plus: piperacillin (100*µg*), nalidixic acid (30*µg*), chloramphenicol (30*µg*), imipenem (10*µg*), cefotaxime (30*µg*), cephalotin (30*µg*), kanamycin (30*µg*), tetracycline (30*µg*), gentamicin (10*µg*), amikacin (30*µg*), ciprofloxacin (5*µg*), fosfomycin (50*µg*), doxycycline (30*µg*), cefepime (30*µg*), ceftazidime (10*µg*), levofloxacin (5*µg*), meropenem (10*µg*), norfloxacin (10*µg*), cefuroxime sod. (30*µg*), tobramycin (10*µg*) ^30^. After 24 hours of incubation at 37°C, the resistance diameters were measured. Clustering of antibiotics and of isolates was performed using the R function heatmap.2 from the R library ^31^ Heatplus and hierarchical clustering of matrices based on Euclidean distances between isolates and between antibiotics.

### Sequencing

To select isolates representative of phenotype, we clustered isolates according to the diameters of inhibition zone against the 20 antibiotics using k-means clustering based on Euclidean distances between isolates (vectors of 20 inhibition zone diameters). The analysis and graphs were produced using R version 3.2.4 ^31^. As clusters may be highly skewed in number of cluster members, we tested all cluster numbers from 1 to 100 and plotted within class sum of squares against *k*. At *k* = 47, the sum of squares tails off and there is a steep local decrease, so that *k* = 47 was fixed as k-means parameter. We obtained 103 isolates, which were subsequently used for sequencing and further analysis. To further validate the choice, we plotted the average number of resistances against number of isolates and antibiotics vs. number of isolates for the total 1178 and the selected 103 isolates (see Supp Fig. 1) and concluded that both distributions are roughly similar. 3000ng DNA were extracted from each of the 103 selected isolates using MasterPure extraction kit (Epicentre) according to the manufacturer’s instructions. Sequencing was performed using Illumina Flex GL.

### Assembly

Genomes were assembled with Abyss (version 1.5.2) ^32^. In order to optimize *k* for the best assembly, k-mer values had to be empirically selected from the range of 20-48 (see Supp. Fig. 2) on a per sample basis to maximize contiguity ^3^. To determine the k-mer length that achieved highest contiguity, the 28 assemblies per draft genome/isolate were compared based on *N50* values. 11 assemblies with an *N50* statistic of less than 5 *×* 10^4^ bp were excluded ^33^.

### Genes

Reference gene clusters were computed from 58 complete *Escherichia coli* genomes (see Table 2) available in June 2015 from NCBI. Genes were identified in wastewater and reference genomes using Prokka (version 1.11) ^34^. Genes were clustered at 80% using CD-HIT ^35^ (version 4.6.3, arguments -n 4 -c 0.8 -G 1 -aL 0.8 –aS 0.8 -B 1). Genes with over 90% sequence identity, but only 30% coverage, as well as genes with 80% or greater identity and covered to phage and virus sequences ^36^ were discarded. A gene cluster is defined to be present in an isolate if there is a Prokka gene in the genome, which is longer than 100 amino acids and has over 80% sequence identity and coverage against the gene cluster representative.

### Pan- and core-genome

To generate the pan- and core-genome size graph we followed the procedure in ^3,16^. We had 92 genomes available. We varied *i* from one to 92. At each subset size *i*, we randomly selected *i* genomes and computed the sizes of the union (pan) and intersection (core) of gene clusters. This random selection was carried out 2000 times in each step.

### Gene clusters to rank genes by correlation to phenotype

Prokka genes were identified in all isolate genomes and then clustered with CD-HIT at 60% sequence identity and 50% coverage (arguments -n 4 -c 0.6 -G 1 -aL 0.8 -aS 0.5 -B 1). A 80% identity cutoff was also tried but dismissed, because the 60% threshold yielded 25% less clusters while adequately clustering homologous gene sequences with lower sequence similarity. This threshold value is also supported by the widespread default use of the BLOSUM62 matrix, the basis of which is sequences clustered by 62% sequence identity.

### Tree

The phylogenetic tree of 92 isolates was built following the procedure of ^37,38^using FastTree version 2.1 ^39^. Sequence reads were aligned to *Escherichia coli* K12 MG 1665 and single nucleotide variant calling was carried out using GATK ^40^. Quality control for variant calling was performed; variants supported by more than ten reads or likelihood score greater than 200 were always in the range of 84 – 99% of variants called per isolate with the exception of 2 isolates where only 59% and 60% of the variants were above the threshold for quality and supporting reads. FastTree 2.1 ^39^ was then used to build the maximum likelihood tree based on the sequences derived from variant calling.

### Phylogrouping

For phylogrouping, the classification system established by Clermont *et al*. ^21^ based on the genes chuA and yjaA and the DNA fragment TspE4.C2 was used. Blast was performed to check each genome assembly for presence or absence of the aforementioned elements with an identity cutoff *≥* 90%.

### MLST

Concerning epidemiology and Multi-Locus Sequence Typing, we used the webserver at https://cge.cbs.dtu.dk/services/MLST/ that follows the MLST scheme in ^41^ for predicting MLSTs from whole genome sequence data ^42^. 92 Draft genome assemblies were submitted and results were obtained; 5 isolates were unidentified demonstrating novel sequence types.

### Virulence factors

Virulence factors protein sequences were downloaded from VFDB: Virulence Factors database ^19,43^. 2000 sequences, which are *Escherichia coli* related, were chosen. Sequences were then clustered at 80% sequence identity using CD-HIT (version 4.6.3, arguments -n 4 -c 0.8 -G 1 -aL 0.8 -aS 0.8 –B 1). A virulence factor was considered present in an isolate’s genome if there is a Prokka gene in the genome that has over 80% sequence identity and coverage against the virulence factor cluster representative.

### ExPEC classification

There are intra- and extra-intestinal pathogenic *Escherichia coli*, which can be classified from the presence of virulence factors ^44-47^. InPEC are characterised by the virulence factors stx1, stx2, escV, and bfpB. They are ExPEC if they contain over 20 of 58 virulence factors afa/draBC, bmaE, gafD, iha cds, mat, papEF, papGII, III, sfa/foc, etsB, etsC, sitD ep, sitD ch, cvaC MPIII, colV MPIX, eitA, eitC, iss, neuC, kpsMTII, ompA, ompT, traT, hlyF, GimB, malX, puvA, yqi, stx1, stx2, escV, bfp, feob, aatA, csgA, fimC, focG, nfaE, papAH, papC, sfaS, tsh, chuA, fyuA, ireA, iroN, irp2, iucD, iutA, sitA, astA, cnf1, sat, vat, hlyA, hlyC, ibeA, tia, and pic.

## Data availability statement

Genome assemblies of the analyzed isolates that support the findings of the study will be made available on the NCBI upon paper publication.

